# Dynamic phase separation of the androgen receptor and its coactivators to regulate gene expression

**DOI:** 10.1101/2021.03.27.437301

**Authors:** Fan Zhang, Samantha Wong, Joseph Lee, Shreyas Lingadahalli, Christopher Wells, Neetu Saxena, Christophe Sanchez, Bei Sun, Ana Karla Parra-Nuñez, Novia Chan, Jennifer M. Bui, Yuzhuo Wang, Paul S. Rennie, Nathan Lack, Artem Cherkasov, Martin Gleave, Jörg Gsponer, Nada Lallous

**Affiliations:** Vancouver Prostate Centre, Department of Urologic Sciences, University of British Columbia, 2660 Oak St., Vancouver, BC, V6H 3Z6, Canada; Michael Smith Laboratories, Department of Biochemistry and Molecular Biology, University of British Columbia, Vancouver, BC, Canada, V6T 1Z4; School of Medicine, Koç University, Rumelifeneri Yolu, Istanbul, 34450, Turkey; Koç University Research Centre for Translational Medicine (KUTTAM), Koç University, Rumelifeneri Yolu, Istanbul, 34450 Turkey

**Keywords:** Androgen receptor, phase separation, transcriptional regulation, prostate cancer

## Abstract

Numerous cancers, including prostate cancer (PCa), are addicted to transcription programs driven by superenhancers (SEs). The transcription of genes at SEs is enabled by the formation of phase-separated condensates by transcription factors and co-activators with intrinsically disordered regions. The androgen receptor (AR), main oncogenic driver in PCa, contains large disordered regions and is co-recruited with the co-activator MED1 to SEs to promote oncogenic programs. In this work, we show that dynamic AR-rich, liquid-like foci form in PCa models upon androgen stimulation and correlate with AR transcriptional activity. The co-activator MED1 plays an essential role in the formation of AR foci while AR antagonists hinder their formation. These results suggest that enhanced compartmentalization of AR and co-activators at SEs may play an important role in the activation of oncogenic transcription programs in PCa. A better understanding of the assembly and the regulation of these AR-rich compartments may provide novel therapeutic options.

## Introduction

Prostate cancer (PCa) is one of the most common cancers in men. Androgen deprivation therapy (ADT) remains the mainstay of systemic therapy for advanced disease, but over time progression to castration-resistant PCa (CRPC) is inevitable^1,2,3^. The androgen receptor (AR) is a ligand-activated transcription factor that drives oncogenic signaling in both castrate-sensitive and - resistant PCa. The transcriptional activity of the AR is tightly regulated. Androgen binding in the cytoplasm leads to translocation of the autoinhibited AR into the nucleus, where it dimerizes and binds to specific androgen response binding sites (ARBSs) to induce transcription of target genes (reviewed in^4^).

Interactions with co-activators such as MED1 are also required for AR transcriptional activity. Increased interactions with and dysregulation of such co-activators have been associated with metastatic and castrate resistant PCa. In addition, amplification of MED1 may promote prostate oncogenesis^5,6^. MED1 has recently been reported to engage AR in androgen-dependent PCa cells at super-enhancer (SE) sites — clusters of enhancers that cover larger genomic areas and present an enrichment of phosphorylated MED1, RNA POLII, p300 and H3K27ac to cooperatively assemble the transcriptional apparatus at high density and induce robust expression of genes that play key roles in defining cell identity^7,8^. The interaction of MED1 and AR at SEs is dependent on CDK7-mediated phosphorylation of the former. The inhibition of MED1 phosphorylation via a CDK7-specific inhibitor reduces AR-dependent tumor growth.

Studies over the last decade have defined abnormal SE-driven transcription programs in cancer that may mark addictions that can be therapeutically exploited^9,10^. SEs contain clusters of transcription factors and co-activators such as MED1 and BRD4 which have both been shown to form nuclear foci at SEs with properties of liquid-like condensates such as dynamic and multivalent interactions among molecules and dependence on environmental changes. Moreover, intrinsically disordered regions in MED1 and BRD4 can form phase separated droplets *in vitro* that can concentrate the transcription apparatus from nuclear extracts, suggesting that SE condensates may ensure robust transcription of cell-identity defining genes in normal as well as cancer cells^7,11,12^.

Given these recent findings, we propose that AR, both a critical differentiation and oncogenic transcription factor, has the ability to form or be recruited to biomolecular foci as mean of transcriptional regulation. Using a variety of molecular biology and biophysical approaches we found that, upon androgen stimulation, full length AR reversibly forms foci that are dependent on the coactivator MED1 and are reduced in the presence of hexanediol, a chemical reported to dissolve proteinaceous condensates. AR transcriptional activity correlates with foci formation and can be modulated by changing cellular foci content genetically or chemically.

## Results

### AR-rich nuclear foci are formed in prostate cancer models upon androgen stimulation

Recent reports suggested that 65% of transcriptionally active H3K27ac SEs in VCaP prostate cancer cells are co-enriched in AR and MED1 upon DHT stimulation compared to only 11% of enhancer regions^13^. Together with the observation that MED1 can form foci at SEs that exhibit properties of liquid-like condensates, these data suggest that the AR may be recruited to transcriptionally active condensates in androgen-positive PCa cells. To investigate this hypothesis, we probed the ability of AR to form foci upon androgen stimulation in PCa cells. We first evaluated whether AR tagged with a non-dimerizing Enhanced Green Fluorescent Protein (AR-mEGFP) forms foci in the prostate cancer LNCaP and LAPC4 cell lines and the normal prostate epithelial RWPE1cell line. Two hours post androgen stimulation (Dihydrotesterone (DHT), 1 nM). AR-rich foci were seen in the nucleus of LAPC4 and LNCaP cells (Fig. 1A). The empty vector expressing the mEGFP protein itself did not show condensate formation in presence or absence of androgens (Figure S1A). Similarly, no foci were observed in the normal prostate epithelial cell line RWPE-1, an AR negative cell line that expresses MED1^13^ (Figure 1A, S1B). We also confirmed that endogenous AR forms foci in these PCa cell lines upon DHT stimulation using immunofluorescence (IF) staining with anti-AR antibody (Fig. 1A). Endogenous AR also formed foci in patient derived xenograft (PDX) PCa tissues (Figure S1C).

**Figure 1:**
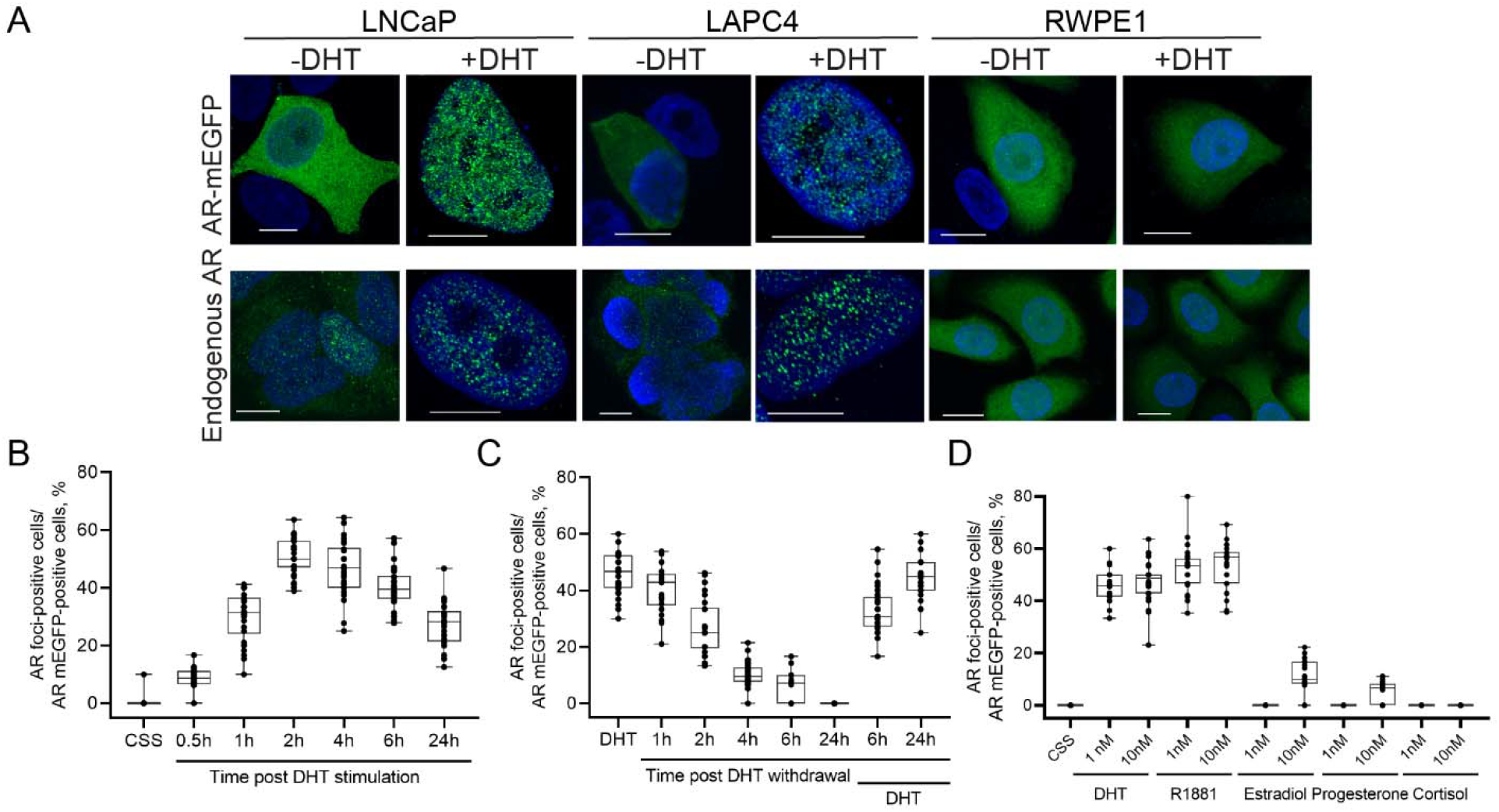
AR-rich condensates form upon androgen stimulation. **A-** Cells were cultured with 5% CSS-containing medium (LNCaP, LAPC4) or K-SFM medium (RWPE-1) for two days for hormone- and growth factor-deprivation and then stimulated with DHT (1 nM) for two h. The mEGFP-AR protein was directly inspected under confocal microscope. The endogenous AR was immunofluorescent (IF) stained with anti-AR antibody and the Alexa Fluor 488-labeld secondary antibody before the confocal microscopy inspection. All scale bar: 10 µm. **B-** LNCaP cells expressing AR-mEGFP were stimulated with DHT (1nM) for various time as indicated. AR-rich condensates were quantified under confocal microscope. The percentages of condensate-containing cells against the AR-mEGFP positive cells were presented. **C-** LNCaP cell expressing AR-mEGFP protein were treated with DHT for 2 h and then the cells were washed twice with PBS to remove DHT. Cells were then cultured in 5% CSS for the indicated time course. DHT was applied back to cells at 6 h and 24 h post DHT removal, respectively. AR-rich condensates were quantified as stated before. **D-** LNCaP cell expressing AR-mEGFP protein were treated with various hormones and condensate formation was quantified. The dotted blot presents the data from at least 30 cells from three independent experiments.

To investigate the reversibility of foci formation, we followed AR-rich foci by a time-lapse live imaging assay in LNCaP cells. Foci containing AR-mEGFP started to form at 30-minutes (min) post DHT stimulation, consistent with the time previously reported for AR nuclear translocation^14^ (Figure 1B, S1D). A similar result was obtained with the synthetic androgen R1881 (Figure S1E). The number of cells with foci peaked at about two hours (Figure 1B, S1D), and dropped rapidly after DHT was removed from the medium, with nearly no foci left after 24 h. However, if the DHT was added back to the media, the foci formed once again (Fig. 1C), demonstrating that AR foci reversibly form upon androgen stimulation. We also investigated if other steroid hormones can stimulate AR foci formation in LNCaP cells. AR-rich foci were specifically induced by 1 nM of DHT and R1881 but only very weakly by high concentrations of estradiol or progesterone (Figure 1D). This specificity in foci formation is consistent with the androgen dependence of AR transcriptional activity that we reported previously^15^. Taken together, our findings demonstrate that AR can form foci in androgen-dependent PCa cells specifically upon androgen stimulation.

### AR-rich foci exhibit properties of liquid-like condensates

Next, we investigated whether AR foci have characteristics consistent with liquid-like phase condensates. A hallmark of condensates are rapid internal reorganization and exchanges with the surroundings, which can be probed via fluorescence recovery after photobleaching (FRAP)^7,16,17^. With this approach we observed that AR-mEGFP foci recovered fluorescence after photobleaching on a time scale of seconds with an average half time of t = 7.7 ± 1.1 s and a mobile fraction of 62% ± 24 (n=21 foci from 15 cells) (Figures 2A-B and S2A). These results are consistent with FRAP measurements collected previously for MED1 and BRD4 foci in embryonic stem cells^7^. However, some of the condensates did not fully recover their fluorescence in the measured time frame, thus the overall 38% of calculated immobile fraction. The percentage of cells presenting AR-rich foci increases quickly with the transfected amount of AR-mEGFP plamid and plateaued at 1 μg, at which the nuclear AR-mEGFP concentration corresponds to 5.9 ± 0.7μM (Figure 2C and S2B). This behavior is reminiscent of the phase behavior that has been previously been measured for G3BP1 *in vitro*^18^. We then tested the effect of 1,6-hexandiol, a compound known to disrupt liquid-like condensates^7,16^, on the formation of AR-rich focus in AR-mEGFP expressing-LNCaP cells. Following HD treatment (4%, 5min), the percentage of foci rich cells reduced by about 50% but AR protein levels remained unchanged (Figure 2D). Moreover, the initial number of cells displaying foci was not recovered 4h after the removal of HD despite the fresh addition of DHT (Figure S2C). Another key feature of condensates is that the saturation concentration for their formation is temperature dependent^17^. To investigate if AR foci are responsive to temperature changes, we stimulated AR-Megfp transfected LNCaP cells with DHT and incubated them under various temperatures for 1h, and then fixed them for quantification by confocal microscopy. As shown in Figure 2E, foci formation was maximal at 37 °C and decreased significantly at lower and higher temperatures with no significant effect on AR protein levels. Overall, these results indicate that AR foci display features of liquid-like condensates.

**Figure 2.**
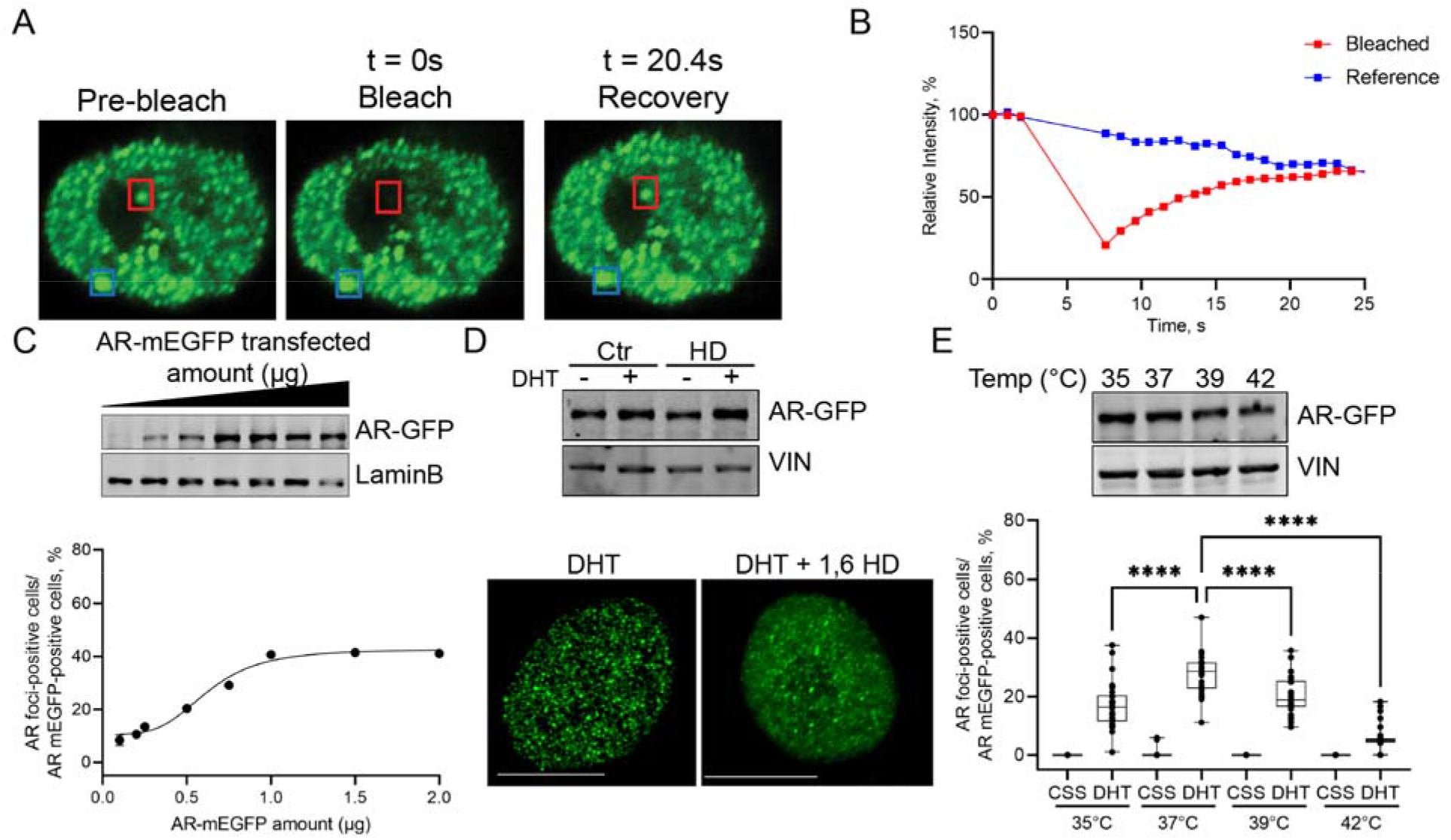
AR-rich condensates present LLPS characteristics. **A-B**-FRAP assay to examine the shuffling of AR in and out the condensates. The AR-mEGFP expressing-LNCaP cells were stimulated with DHT for 2 h. The condensate was photobleached with 100% of laser power for 10 seconds, the time-lapse imaging on the bleached punctum as well as a reference punctum that was not bleached were captured with confocal microscope in a Z-stacking model. The steal images before, on and after the photobleaching were shown (A). The red frame indicated the bleached spot and the blue frame showed the unbleached reference spot. A representative image showing the relative fluorescence intensity of the two regions is presented (B). **C-** LNCaP cells transfected with increasing amount of AR-mEGFP plasmid were cultured in 5% CSS media for 2 days. Cells were then stimulated with DHT for 2h and AR-rich condensates were quantified under confocal microscope. The percentages of condensate-containing cells against the AR-mEGFP positive cells were presented. Values are expressed as mean ± SEM from 3 independent experiments. The corresponding nuclear AR levels were evaluated by Western bot by using GFP antibody **D-** AR-mEGFP expressing-LNCaP cells were stimulated with DHT for 2 h and then received 4% 1,6-HD for 5 min. Cells were then fixed for confocal microscopy imaging. The effect of HD on AR protein level was evaluated by western blot by using anti-GFP antibody. **E-** The AR-mEGFP expressing-LNCaP cells starved in 5% CSS for two days were cultured in incubators with various temperatures for 1 h with DHT. Cells were fixed and AR-rich condensates were quantified under confocal microscope. The dotted blot presents the data from at least 30 cells from three independent experiments. p values are indicated by stars **** < 0.0001. The dotted blot presents the data from 30 cells from three independent experiments. The effect of temperatures on the stability of AR protein was evaluated by western blot by using GFP antibody.

### AR foci formation and transcriptional activity correlate in PCa cells

To correlate AR foci with AR transcriptional activity, we first studied the colocalization of AR-rich foci and nascent RNA (Figure 3A, S3A). AR-mEGFP transfected LNCaP cells were stimulated with DHT for 2h, labelled with 5-Bromouridine-5′-triphosphate, fixed and stained. The newly synthesized RNA was visualized using an anti-Bromodeoxyuridine (BrdU) antibody. As seen in Figure 3A (top), the majority of AR foci colocalized with nascent RNA. A similar level of colocalization was found with endogeneous AR (Figure S3A). MED1 interacts with AR and facilitates the recruitment of RNA polymerase II (Pol II) and the initiation of transcription^13,19^. Androgen stimulation in PCa cells induces MED1 phosphorylation at threonine T1457, which enables MED1’s physical interaction with AR at SEs^13^. To characterize the role of phosphorylation on foci formation, we tested colocalization of AR foci with phosphorylated MED1 (pMED1, at T1457) and phosphorylated Pol II (pPol II, at S2). Using phospho-specific antibodies we observed that a significant number of the AR containing foci colocalized with pMED1 and pPol II in LNCaP cells overexpressing AR-mEGFP (Figure 3A) or stained for endogenous AR (Figure S3A). To further validate the findings, we performed proximity ligation assay (PLA) to detect the interaction between endogenous AR, pMED1 and pPol II, respectively, as well as the colocalization of their interaction sites with AR-mEGFP containing foci (Figure 3B, S3B). These experiments clearly demonstrated that AR, pMED1 and pPol II interact, and that they are co-present in many, but not all, AR foci.

**Figure 3:**
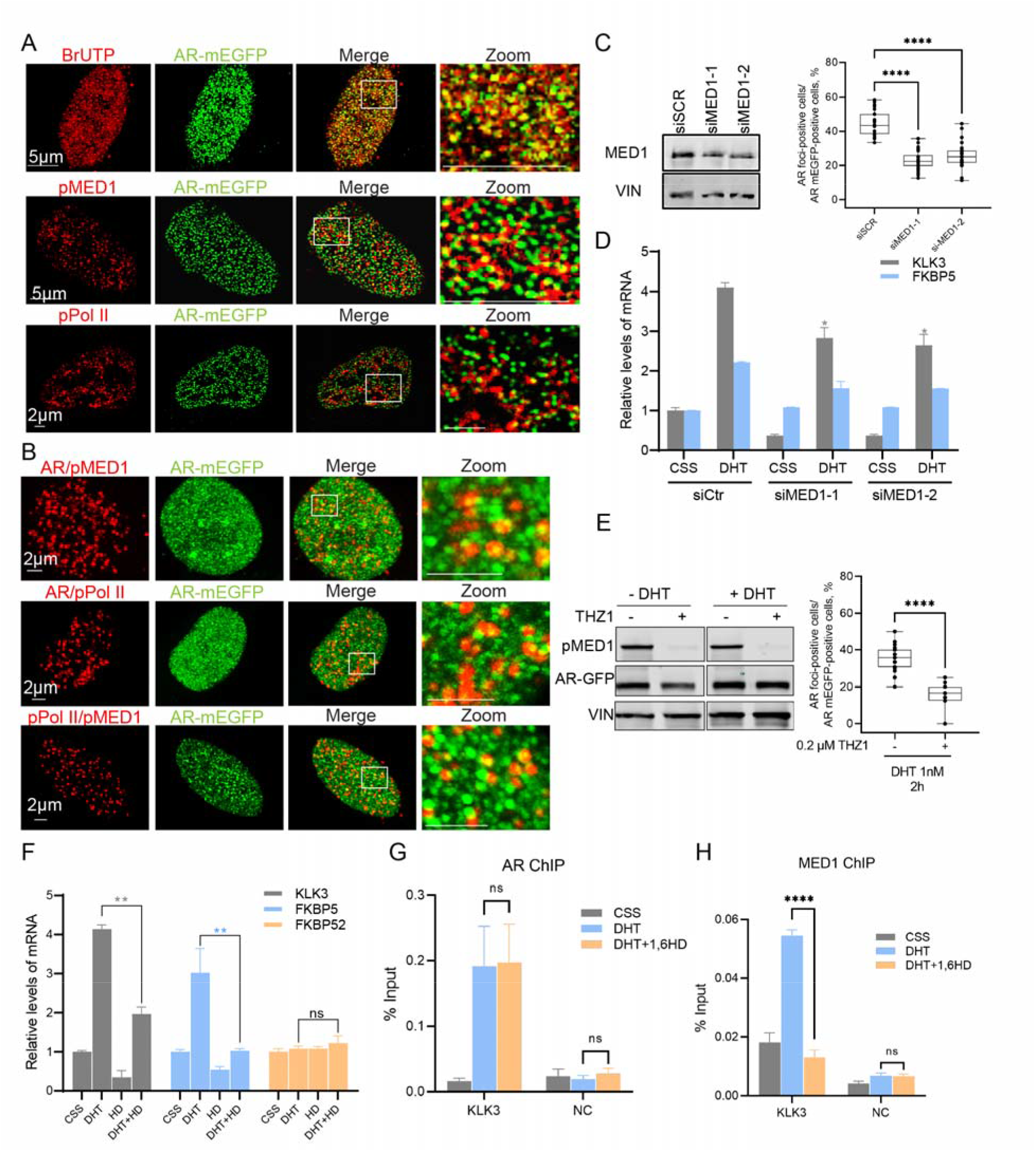
Condensate formation correlates with AR transcriptional activity. **A-** LNCaP cells starved with 5% CSS for two days were stimulated with DHT for 2 h. IF on BrUTP, pMED1 and pPol II were performed (red panels) and the co-localization with AR-rich condensates (green panel) were examined under confocal microscope. **B-** Combined PLA staining (in red) and AR-mEGFP (green panel) were carried out in DHT-stimulated LNCaP cells. The colocalization of PLA signal with AR-rich condensates was inspected under the confocal microscope. Images in the white frames were enlarged and displayed in the right panel. **C-D** MED1 was knocked down in LNCaP cells and the levels of MED1 proteins were validated with western blot. Vinculin (VIN) was used as a loading control. Effects of knocking down of MED1 and AR condensates (C) and on AR transactivation (D) were evaluated. The PCR values are expressed as mean ± SD. The dotted blot presents the data from at least 30 cells from three independent experiments **E-** Inhibiting MED1 phosphorylation by THZ1 (0.2 µM for 2h) in AR-mEGFP expressing LNCaP cells, reduces foci formation without significantly affecting AR protein levels as shown by western blot **F-** The mEGFP-AR expressing-LNCaP cells were treated with DHT with / without 2.5% 1,6-hexanediol (HD) for 30min, and then cells were washed with PBS twice and incubated with DHT-containing medium for 16 h. Total RNAs were extracted and the mRNA levels of genes of interested were examined using qRT-PCR. Values are expressed as mean ± SD. **G-H** LNCaP cells were hormone starved in 5% CSS for 72h then treated with 1nM DHT or ETOH for 2h hours and followed by 2.5% 1,6 HD for 30 mins. The crosslinked chromatin was incubated with AR (G) or MED1 (H) specific antibodies. Enrichment of AR and MED1 at KLK3 site were quantified by qPCR using specific primers. p values are indicated by stars: ns ≥ 0.05, * 0.01 to 0.05, ** 0.001 to 0.01, *** 0.0001 to 0.001, **** < 0.0001.

Next, we investigated the impact of MED1 repression on foci formation and transcriptional activity of AR. Knocking down the expression with siMED1 significantly reduced AR-rich foci formation (Fig. 3C). The effect of MED1 silencing on foci formation was mirrored by a lower AR transcriptional activity as seen by the reduced mRNA levels of AR-targeted genes *KLK3* and *FKBP5* (Fig. 3D), two genes that have been found within 100kb of SE peaks in PCA cells ^13^. We also found that the inhibition of MED1 phosphorylation by using the CDK7 inhibitor THZ1, previously shown to reduce AR recruitment to SE^13^, reduces foci formation (Fig 3E). Taken together, our analyses suggest that AR can form phase-separated condensates at SEs in AR^+^ PCa cells. If so, HD treatment, which can dissolve coactivator condensates, should impact transcriptional activity and occupancy of SE-driven genes. Indeed, mRNA levels of AR-targeted genes *KLK3* and *FKBP5* were significantly reduced upon HD treatment (shown to reduce AR-rich condensate in Figure 2D) without any off-target effect on non-AR regulated gene FKBP52 (Figure 3F). To characterize this mechanism, we conducted chromatin immunoprecipitation (ChIP) of AR (Fig 3G) and MED1 (Fig 3H) in presence and absence of HD. We observed a strong enrichment of AR and MED1 binding following androgen stimulation at the *KLK3* SE (Figure 3G-H). Interestingly, HD treatment did not affect AR binding to chromatin, however it significantly reduced MED1 binding to AR SE to a near basal level. Overall, these findings demonstrate that reducing AR foci formation genetically or chemically represses AR-target gene expression but not DNA binding in LNCaP cells. This suggest that some foci are coactivator condensates to which AR partitions.

### The full-length AR is required for foci formation

Biomolecular condensate formation is a common property of ID activation domains of transcription factors and coactivators such as MED1^7,12^. However, recent studies revealed that the essential property of scaffolds that drive the formation of phase separated stress granules is their multi-domain (multivalency) character^20^. In order to identify, which AR domain(s) are necessary for foci formation in PCa cells, we generated mEGFP-tagged truncations and tested foci formation upon DHT stimulation and validated that all proteins were expressed to similar levels (Figure 4A and S4A). Interestingly, foci were only seen in presence of full length AR after DHT stimulation and not with any of the truncated domains (Fig. 4B). Surprisingly, even the constitutively active splice variant V7, which lacks the LBD, did not form foci, suggesting an important role of the LBD in this mechanism. Together these results suggest that the NTD, DBD and LBD are required for foci formation. It is important to note that the ability to form nuclear foci depends on the different constructs’ ability to localize to the nucleus. However, all constructs were able to localize to the nucleus, the only exception being the construct that contains the LBD only, which is consistent with previous reports^21,22^.

**Figure 4:**
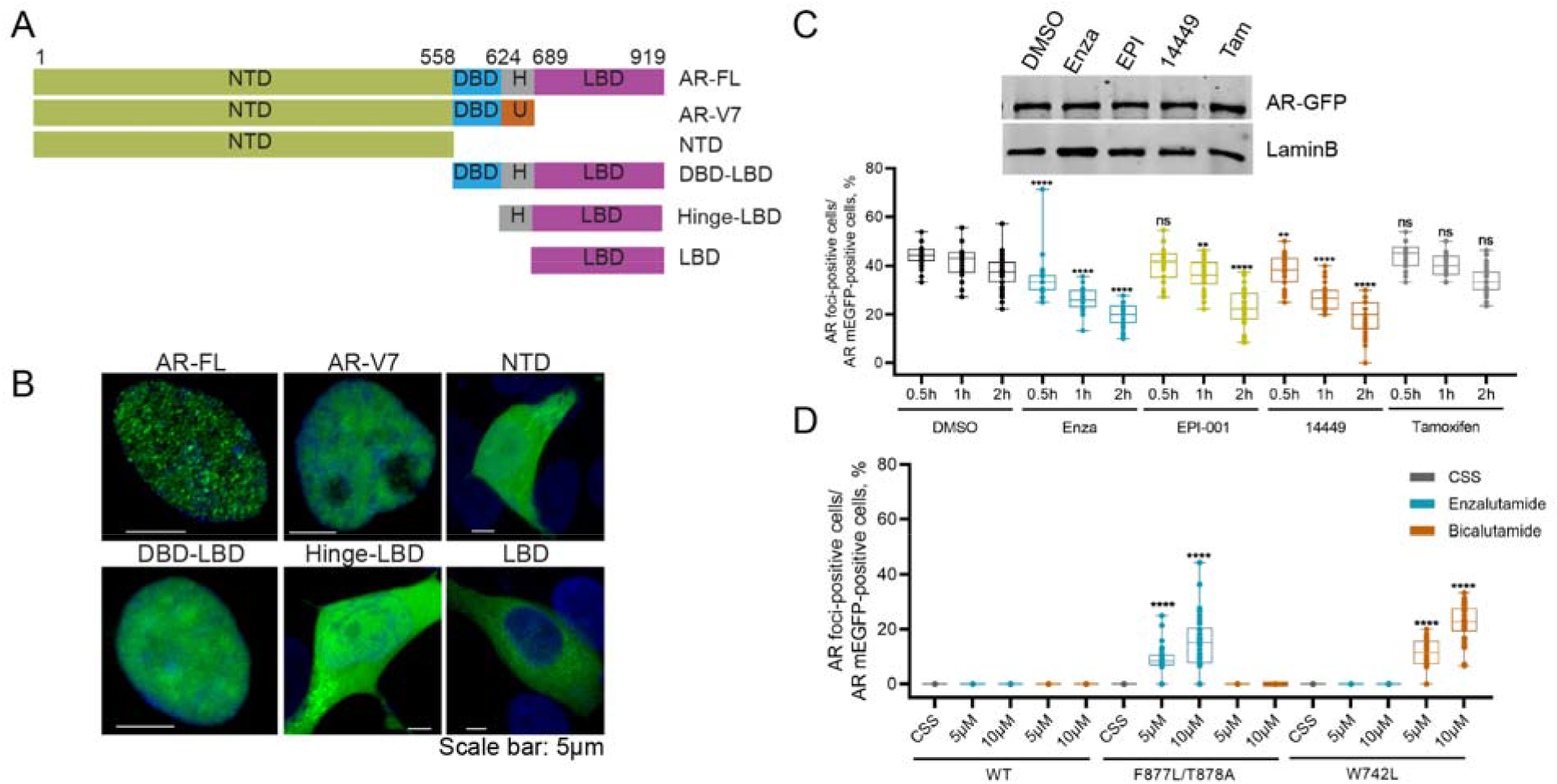
Full Length AR is required for condensate formation. A-Truncated forms of AR-mEGFP. N-terminal domain (NTD), DNA-binding domain (DBD), hinge region (H) and the ligand-binding domain (LBD). AR variant protein V7 presents its unique cryptic exon inclusion sequence (U). **B-** Condensate formation by the mEGFP tagged truncated AR forms was evaluated in LNCaP cells using confocal microscopy. **C-** AR-mEGFP transfected-LNCaP cells were stimulated with DHT for 2 h and then received treatment with 10 µM of enzalutamide (Enza), EPI, 14449 and Tamoxifen (Tam) for 30 min, 1h and 2h. The puncta were quantified under confocal microscope. Western blot showing nuclear AR levels 2h post treatment with the studied inhibitors **D-** Plasmids expressing mEGFP-tagged wild type (wt)-AR as well as F877L/T878A and W742L AR mutants were transfected into LNCaP cells. Cells were starved with 5% CSS for two days and then increasing concentrations of bicalutamide or enzalutamide were added for 2h. Condensates were then quantified. The dotted blot presents the data from at least 30 cells from three independent experiments.

To better understand this mechanism, we tested the effect of AR antagonists that target the NTD (EPI-001)^23^, DBD (VPC-14449)^24^ or LBD (Enzalutamide, Enza)^25^ on foci formation. In this work, the ER antagonist (tamoxifen) was included as a negative control. AR-mEGFP transfected LNCaP cells were stimulated with DHT for 2h to ensure AR nuclear translocation and foci formation, and then treated with 10µM of the tested drugs for an additional 2h. Similar to our truncation experiments, foci formation was significantly reduced 30 minutes post treatment with enzalutamide and 14449 and 1h post treatment with EPI-001, with no significant effect on the nuclear levels of AR (Figure 4C); as expected, tamoxifen had no significant effect on AR-rich foci. Similar effects were seen when LNCaP cells were treated with AR antagonists prior to androgen treatment, except for enzalutamide that inhibited foci formation more significantly (Figure S4B). This additional effect could be attributed to reduction of AR nuclear translocation for enzalutamide^3^. Lastly, we evaluated the ability of two AR-LBD mutants F877L/T878A^15^ and W742L^26^, reported to respectively convert enzalutamide and bicalutamide into agonists, to form foci in the absence of androgens. The mEGFP-tagged mutants as well as the wild type (WT)-AR plasmids were transfected into CSS starved LNCaP cells. As expected, no foci were formed with the wild-type AR in the absence of androgen stimulation. However, enzalutamide and bicalutamide induced F877L/T878A-rich and W742L-rich foci, respectively, in a dose dependent manner (Figures 4D and S4C) which is concordant with previously reported transcriptional activity associated with those mutants^15^. Overall, these experiments demonstrate that the multi-domain character of AR is key to its ability to form foci in PCa cells.

## Discussion

Transcriptional coactivators such as BRD4 and MED1 have been shown to form nuclear foci at SEs that exhibit properties of liquid-like condensates. Moreover, the intrinsically disordered C-terminus of MED1 forms phase-separated droplets that can concentrate the transcriptional machinery from nuclear extracts. Here, we show that AR, a known interactor of BRD4 and MED1^27,28^, also form nuclear foci upon specific stimulation with androgens (Figure 1) in AR^+^ PCa cells. The AR-rich foci exhibit properties found in liquid-like condensates such as responsiveness to HD treatment or temperature changes and recovery of fluorescence after photobleaching (Figure 2). Importantly, the recovery half live calculated from FRAP experiments on AR foci is similar to the one previously measured for MED1 foci, indicating a colocalization in same foci upon DHT stimulation (Figure 3). We also demonstrate that foci formation is dependent on the multi-domain character (multi-valency) of AR, a characteristic feature of phase separation scaffolds. Truncated receptor forms lacking the NTD, the DBD or the LBD are not sufficient to induce foci despite being expressed to similar levels as full length AR and being able to translocate to the nucleus. Similarly inhibitors of the DBD, LBD and NTD, required for DNA binding, androgen ligation, transactivation and co-factor binding, respectively, each reduced foci formation (Figure 4). Taken together, these results suggest that AR foci could be phase-separated condensates. However, recent studies suggest that similar features can also be seen in transiently formed compartments through processes distinct from phase separation. Indeed, it has been shown that replication foci formed during lytic Herpes Simplex Virus infection have some of the same properties as the AR foci, but that they are not formed via phase separation^29^. In this context, it is interesting to note that not all AR foci displayed substantial recovery after photobleaching and that HD treatment reduced the number of cells with foci only by about 50%. Thus, more studies are required to fully capture the biophysical properties and the assembly mechanisms of AR foci.

Independent on whether AR foci induced in PCa cells are all truly liquid-like condensates, our results reveal that these compartments are sites of transcriptional activity. Essentially all AR foci co-localize with newly synthesized RNA in LNCaP cells and a significant fraction does so with phosphorylated MED1 (T1457) and RNA Pol II as seen by immunofluorescence and PLA assays (Figure 3). This finding suggests that androgen-induced AR foci are transcriptional hubs in AR^+^ PCa cells. Reduced expression of one of the biomolecular partners of AR in these hubs, MED1, lowered the number of cells with foci and AR transcriptional activity to a similar extend (Figure 3 C-D). Our findings are of particular interest in light of a recent study demonstrating that phosphorylation of MED1 at T1457 upon androgen stimulation is essential for AR-mediated transcription in androgen-positive PCa cells, i.e., the transcription of AR targets genes (e.g..*KLK3, TMPRSS2*) and oncogenic drivers (e.g. *ERG* and *MYC)*. Rasool *et al*. also revealed that MED1 phosphorylation can be impeded by a CDK7-specific inhibitor, THZ1, which blocks genome-wide co-recruitment of AR and MED1. Consistent with this result, we find that THZ1 reduces AR foci formation^13^ (Figure 3E). Together these findings suggest that nuclear AR proteins can partition coactivator at SEs and induce foci with liquid-like properties. We found that HD, which can lower foci numbers and SE occupancy of coactivators in embryonic stem cells, also significantly reduces AR foci numbers and the mRNA levels of two AR target genes located within 100kb of SE peaks in PCa cells. Interestingly, inhibition of foci formation by HD disrupted MED1 recruitment to AR DNA binding sites, but did not affect AR binding to its target gene. This result points toward a partitioning of AR to coactivator condensates as a downstream event of SE binding by the AR (Figure 2 and 3).

Under physiological conditions, SE programs control genes involved in cell. In cancer cells, by contrast, oncogenes acquire SE in order to assemble the transcriptional apparatus at high density and enable enhanced transcription of target genes^7,10,30^. Mechanisms controlling the SE acquisition by oncogenic drivers include enhancer amplification, enhancer hijacking, small insertions that create master transcription binding site and oncogene overexpression (reviewed in ^10^). The dependence on SE programs has been reported for many cancers, including PCa^9,13,31^. The majority of advanced PCa, including CRPC, are addicted to AR signaling and drive SE-dependent oncogenic programs such as *TMPRSS2-ERG* and *MYC*. How this addiction is established at molecular level is not fully clear, but increased levels of pMED1 and BRD1 have been reported to be negatively prognostic in men with metastatic PCa^13,27,32^. AR overexpression in CRPC is associated with BRD4 dependent chromatin relaxation^33^, and BRD4 associated SE programs lead to the expression of key oncogenes in CRPC such as TMPRSS2-ERG^9^. In this context, it is interesting to note that we only observed AR foci formation upon androgen stimulation in PCa cells, while they were not detected in prostate normal epithelial cell RWPE-1 despite the transient re-expression of AR and the presence of MED1 in those cells. This suggests that the presence of AR and MED1 is not sufficient to induce foci formation in normal cells and that additional oncogenic programs such MYC or ERG may be required.

In summary, we reveal that dynamically formed foci are important to the transcriptional activity of AR in androgen-sensitive PCa cells. This transcriptional activity can be modulated by changing the foci content genetically or chemically. A better understanding of foci assembly mechanism and their mode of action may open novel therapeutic venues to treat advanced forms PCa, specifically those that are addicted to androgen-driven SE transcriptional programs.

## Methods

### Cell lines

Human cell lines LNCaP, LAPC4 and RWPE-1 were received from the American Type Culture Collection (ATCC). LNCaP and LAPC4 cells were maintained in RPMI-1640 media (Invitrogen Life Technologies, Burlington, ON) containing 10% heat inactivated fetal bovine serum (FBS; Invitrogen Life Technologies). RWPE-1 cells were grown with the Keratinocyte Serum Free Medium (Gibco) with 0.05 mg/ml bovine pituitary extract (BPE) and human recombinant epidermal growth factor (EGF). For the treatment with charcoal-stripped serum (CSS), cells were grown in 5% CSS (Invitrogen Life Technologies) in phenol-red free RPMI-1640 medium (Invitrogen Life Technologies). For starvation in RWPE-1 cells, cells were grown in the base medium without BPE and EGF. All the cell lines were tested negative from mycoplasma contamination and authentication was confirmed by genomic sequencing or STR profiling.

### Antibodies and Reagents

Primary antibodies used in this study were: anti-AR from Santa Cruz Biotechnology (sc-7305); anti-MED1 (A300-793A) from Bethyl; anti-phospho-MED1 (ab60950), anti-phospho-RNA Rol II (ab5408) from Abcam; anti-vinculin from Sigma-Aldrich (V9131); anti-GFP (#2956) from Cell Signaling Technology. Secondary antibodies for western blot were: donkey anti-mouse IgG (IRDye 680CW #926-68072; 800CW, #926-32212) and donkey anti-rabbit IgG (IRDye 680CW, #926-68073; 800CW, #926-32213), all from Li-Cor (LI-COR Biosciences). Secondary antibodies for immunofluorescence (IF) staining were as follows: donkey anti-rabbit Alexa Fluor 594 (A-21207) and donkey anti-mouse Alexa Fluor 488 (A-21202), both from Invitrogen Life Technologies. The dilutions of antibodies were 1:1000 for primary antibodies and 1:3000 for secondary antibodies for western blot; and 1:200 for primary antibodies and 1:500 for secondary antibodies for IF staining. Dihydrotestosterone (DHT) was purchased from Steraloids (Newport, RI, US) and enzalutamide from Haoyuan Chemexpress Co. (Shang Hai, China). β-etradiol, hydrocortisone, progesterone, tamoxifen, bicalutamide, EPI-001, 1,6-Hexanediol (#240117), 5-Bromouridine 5′-triphosphate (BrUTP) were purchased from Sigma Aldrich.

### Constructs

mEGFP-N1 was a gift from Michael Davidson (Addgene plasmid # 54767 ; http://n2t.net/addgene:54767 ; RRID:Addgene_54767). AR constructs were cloned at the Nterminus of the mEGFP tag by using the restriction enzyme sites NheI and BamHI. AR mutants were made by QuikChange II Site-Directed Mutagenesis Kit (Agilent Technologies).

### Transfection

Transfection of plasmids was carried out using Lipofectamin 3000 (Invitrogen Life Technologies) according to the manufacturer’s user guides. 1 μg of plasmid was transfected into one well of the 24-well plate unless otherwise indicated. Transfection of siRNAs was performed using Oligofectamin (Invitrogen Life Technologies) following the user guide. The siRNAs targeting MED1 were purchased from Applied Biological Materials (Abm, Richmond, BC, Canada, Cat# i5129621). The siRNA for the scramble control (5′-CAGCGCUGACAACAGUUUCAU-3′) was from Dharmacon.

### Immunofluorescence staining and confocal microscopy

Cells grown on glass coverslips were fixed with 3.5% paraformaldehyde (PFA) and then permeabilized with 0.5% Triton x-100. After blocking with 3% skim milk, cells were incubated with the primary antibody diluted in 3% milk for 1 hour at 4 °C for overnight followed with 1 hour incubation of the secondary antibody at 37 °C for 40 minutes in dark. Coverslips were then mounted on slides with DAPI-containing mounting medium (VECTOR, Burlingame, CA) and stored at 4 °C in dark until being examined with microscope. Fluorescence staining was visualized with an Olympus FV3000RS confocal microscope using a 60x UPLAPO oil objective. The images were taken with the FV31S-SW software with the confocal pinhole set to “automatic” under the Z-stack model. For presentation purposes, images were exported as bitmap (BMP) files. The condensates quantification was performed by two independent investigators and presented as the percentage of cells containing the condensates to the cells expressing GFP-tagged proteins. The results shown are the average of three independent experiments. The graph was prepared with the GraphPad Prism 9 software.

### Fluorescence recovery after photobleaching (FRAP)

FRAP analysis was performed with a Zeiss confocal laser scanning microscope LSM780 using a Plan-Apochromat 63× 1.40 oil immersion objective. The selected condensates were bleached with the argon laser with 100% of laser power, and the fluorescence recovery was monitored with continuous z-stack scanning for 30 seconds. The images of a non-bleached region were also captured simultaneously from the same field for the reference purpose. A total of 21 foci were monitored for the FRAP assay and the half-lives and mobile fractions of AR-mEGFP-rich condensates were analyzed with Zeiss ZEN software (ZEN2010) and graphed using GraphPad Prism 9 software.

### Proximity ligation assay (PLA)

For the PLA analysis, the protein-protein interaction was identified using a commercially available PLA kit (Duolink; Sigma-Aldrich) as described before ^34^. Briefly, cells were stained with a pair of primary antibodies at 4 °C for overnight and then labeled with probes linking with complementary DNA (cDNA) at 37°C for 1 hour. cDNAs were ligated at 37 °C for 40 minutes and the product was amplified at 37 °C for 100 minutes. The signal was inspected using the Olympus FV3000RS confocal microscope with a 60x UPLAPO oil objective.

### BrUTP incorporation assay

BrUTP was transfected into cells with Lipofectin (Invitrogen Life Technologies) following the user guide. The transfection solution containing 10 mM BrUTP was incubated with cells for 30 min at 37 °C and then replaced with culture medium. Cells were fixed after 2 hour with 3.5% PFA and proceeded with IF staining with anti-BrdU antibody (ab6326, Abcam) followed with the secondary antibody conjugated with Alexa Fluor 594.

### Live cell imaging

Cells grown in glass-bottom dish were cultured in a chamber supplied with 5% CO_2_ at 37 °C with humidity on the FV3000RS confocal microscope. Time lapse live images for one minute interaction were taken using the 60x UPLSAPO silicon oil lens. The Z-stack model and the Z Drift Compensator were controlled with the FV31S-SW software.

### Quantitative reverse transcription PCR

RNA extraction and reverse transcription PCR (RT-PCR) were performed as described previously ^35^. Real-time monitoring of PCR amplification of cDNA was carried out using FastStart Universal SYBR Green Master (Rox, Roche). The primers used for gene expression are: *KLK3*: forward 5′-GGGTGTCTGTGTTATTTGTGG-3′, reverse 5′-TTGCTGTGAGTGTCTGGTG-3′; *FKBP5*: forward 5′-CACCCACTTGAAGTCCTGTATC-3′, reverse 5′-AGGAGCAGCCAGACCTATTA-3′; *FKBP52*: forward 5′-CACTACACTGGCTGGCTATT-3′, reverse 5′-TTCACCGCGAGTCTGTATTC-3′; *Vinculin*: forward 5′-TCCTGTAATTCCTACCTCCCTG-3′, reverse 5′-TGCCCTCTCATTTTGCCTAG-3′. Target gene expression was normalized to the levels of Vinculin in respective samples as an internal control.

### Western blot analysis

Protein (20 µg) was separated electrophoretically on SDS polyacrylamide gels and then transferred to a nitrocellulose membrane (BIO-RAD Laboratories, Mississauga, ON). After blocking, membranes were incubated with a primary antibody at 4 °C overnight followed by incubation with a secondary antibody labeled with fluorescent dye. Protein was visualized using the Odyssey Imaging System (LI-COR Biosciences). Vinculin was included as loading control. Nuclear concentration of AR was estimated by using a standard curve obtained with recombinant AR protein.

### Chromatin immunoprecipitation (ChIP)-PCR

LNCaP cells were hormone starved in phenol red free RPMI media supplemented with 5% CSS for 3 days. Cells were then treated with 1nM DHT or ETOH for 2 hours and followed by the treatment with 2.5% 1,6 hexanediol or PBS for 30 mins. The cells were immediately washed with PBS and cross-linked with 1% formaldehyde for exactly 10 minutes at room temperature. After quenching with 100mM glycine and extensive washing with cold PBS, the cells were harvested. Nucleus enriched cell pellets were obtained by sequential lysis with LB1 (50 mM HEPES-KOH (pH 7.6), 140 mM NaCl, 1 mM EDTA, 10% (vol/vol) glycerol, 0.5% (vol/vol) NP-40/Igepal CA-630 and 0.25% (vol/vol) Triton X-100), LB2 (10 mM Tris-HCl (pH 8.0), 200 mM NaCl, 1 mM EDTA and 0.5 mM EGTA). Finally, the nucleus was lysed in LB3 (10 mM Tris-HCl (pH 8.0), 100 mM NaCl, 1 mM EDTA, 0.5 mM EGTA, 0.1% (wt/vol) sodium deoxycholate and 0.5% (vol/vol) N-lauroylsarcosine) and the chromatin was sheared to 300 – 800 base pair (bp) fragments by Bioruptor (Diagenode, Belgium) 12 cycles of 30 second “ON” and 30 second “OFF” intervals. After separating 5% of the sample as ‘input’, the rest was used for IP. For IP, the crosslinked chromatin was incubated with AR or MED1 conjugated Protein G beads (Dynabeads, Thermo Scientific) overnight with slow rotation at 4°C. The next day, chromatin from both IP and ‘Input’ samples were eluted and de-crosslinked at 65°C for 16 hours followed by DNA isolation by PCR and DNA cleanup kit (NEB). Enrichment of AR and MED1 at specified enhancers were quantified by qPCR using specific primers. The primers used are: *KLK3*: forward 5′-ACAGACCTACTCTGGAGGAAC-3′, reverse 5′-AAGACAGCAACACCTTTTT-3′; *NC*: forward 5′-CATTTCCTGCTTGTCCTCTG-3′, reverse 5′-GGCCTTCCTGGTATGAAATG-3′.

## Author contributions

FZ, JG and NL designed research; FZ, SW, JL SL, CW, NS, CS, BS, AKPN, NC performed research; PSR, YW, NAL, AC, MEG contributed new reagents/analytic tools; FZ,SL, NAL, JMB, JG, and NL analyzed data; and FZ, MEG, NAL, JG, and NL wrote the paper.

