## Supplementary information for "Dynamic phase separation of the androgen receptor and its coactivators to regulate gene expression"

### **Supplementary figures:**

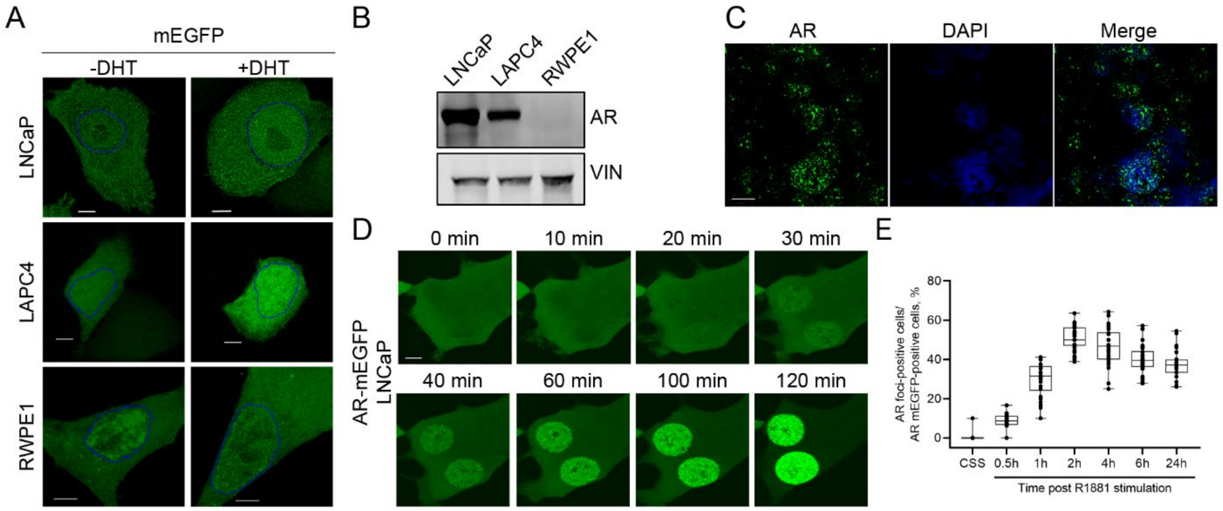


**Figure S1:** AR-rich condensates form upon androgen stimulation. **A-** The vector encoding mEGFP protein doesn't form puncta upon androgen stimulation. mEGFP-transfected cells were starved in 5% CSS-containing medium (LNCaP and, LAPC4) or K-SFM medium (RWPE-1) for two days and then stimulated with 1 nM of DHT for 2 h. The localization and distribution of mEGFP protein was inspected under confocal microscope. Nucleus was outlined in blue. Scale bar: 5 µm. **B-** Western blot showing endogenous AR levels in the studied cells. **C-** The patient derived xenograft (PDX) PCa tumor lines were grafted to mice supplemented with testosterone (10 mg/mouse). OCT blocks of tumor tissue were prepared for the tumor tissues and 5 µm thick tissues were sectioned for. IF was performed on AR. The images were taken with the confocal microscope using the Z-stack model. Scale bar: 10 µm. **D-** The mEGFP-AR protein-transfected LNCaP cells were hormone starved (5% CSS) for 2 days and then stimulated with 1 nM DHT. A live time-lapse confocal imaging assay was performed along the DHT treatment. **E-** LNCaP cells expressing AR-mEGFP were stimulated with R1881 (1nM) for various time as indicated. AR-rich condensates were quantified under confocal microscope. The percentages of condensate-containing cells against the AR-mEGFP positive cells were presented. The dotted blot presents the data from at least 30 cells from three independent experiments.

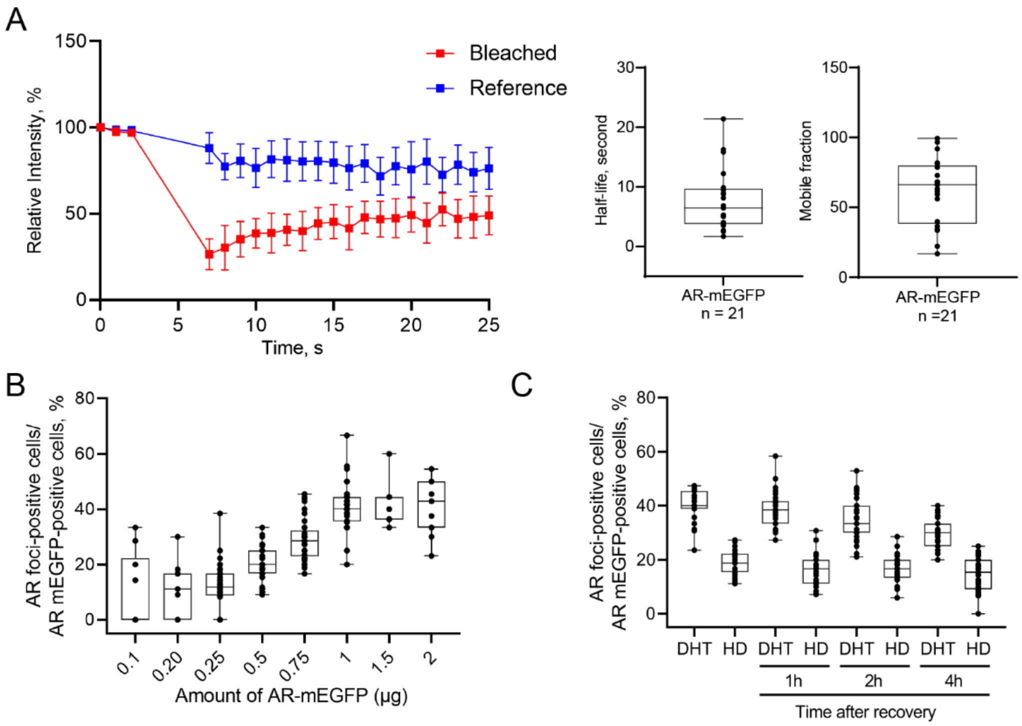


**Figure S2:** AR-rich condensates present LLPS characteristics. **A-** FRAP results are presented as mean ± SD (n =21) and the corresponding quantification of half-life and mobile fraction. **B-** LNCaP cells were transfected with increasing amount of AR-mEGFP plasmid and cultured in 5% CSS for 2 days. Cells were then stimulated with DHT for 2h and AR-rich condensates were quantified under confocal microscope. **C-** The mEGFP-AR expressing-LNCaP cells were stimulated with DHT for 2 h and then received 4% 1,6-HD for 5 min. Cells were then washed with PBS twice and incubated with DHT-containing medium for the indicated time course. AR-rich condensates were then quantified. The dotted blot presents the data from at least 30 cells from three independent experiments.

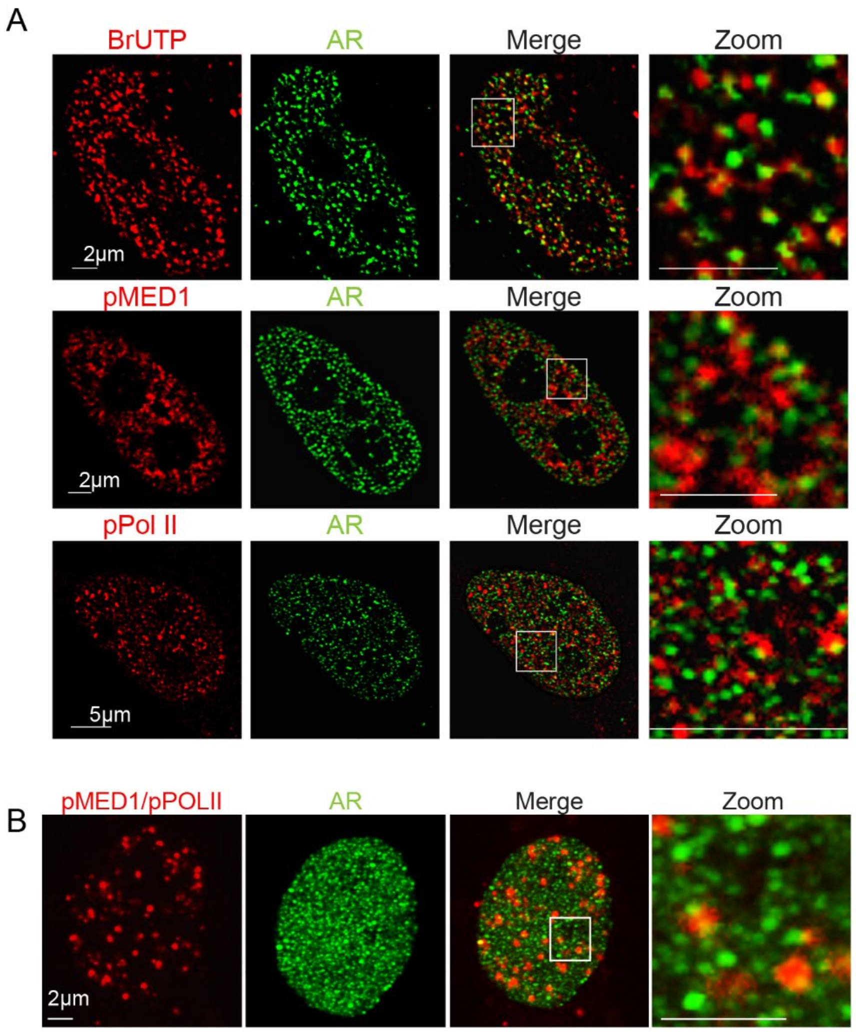


**Figure S3:** Condensate formation correlates with AR transcriptional machinery. **A-** LNCaP cells starved with 5% CSS for two days were stimulated with DHT for 2 h. IF on BrUTP, pMED1 and pPol II (red panels) and endogenous AR (green) were performed. The co-localization with AR-rich condensates was examined under confocal microscope. **B-** Combined PLA staining (in red) and IF staining of AR (green panel) were carried out in DHT-stimulated LNCaP cells. The colocalization of PLA signal with AR-rich condensates was inspected under the confocal microscope. Images in the white frames were enlarged and displayed in the right panel.

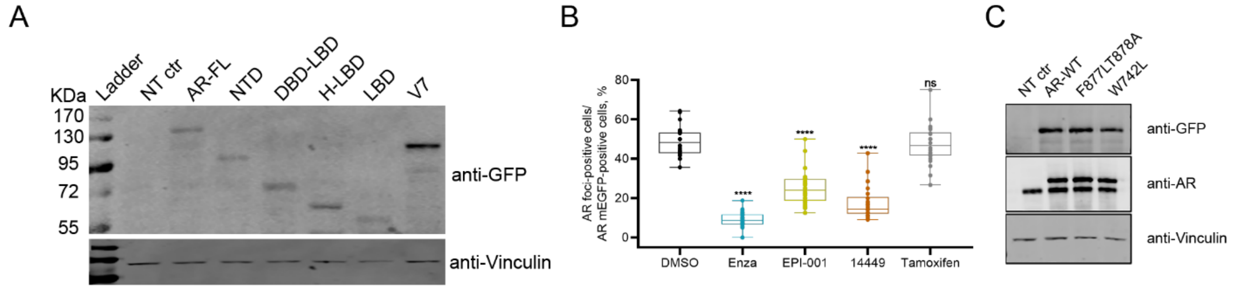


**Figure S4:** Full Length AR is required for condensate formation. **A-** Western blot showing expression levels of mEGFP-tagged AR truncations. **B-** AR-mEGFP transfected LNCaP cells were pre-treated with the compounds for 2 h prior to the DHT stimulation of 2 h. Condensate formation was then quantified. The dotted blot presents the data from at least 30 cells from three independent experiments. **C-** Western blot showing the expression levels of wild-type and mutated forms of AR.

**Supplementary method:**

***Patient-Derived Xenografts (PDX) assay***

The patient-derived prostate tumor tissue lines were implanted in male NOD-SCID mice supplemented with testosterone (10 mg/mouse), as previously described (Dong L, Cancer Research 2014, PMID 24356420). When tumors reached 500 $mm^3$, the animals were sacrificed and tumors were dissected to prepare for the frozen OCT blocks. 5 μm thick tissues were sectioned from the OCT block for the IF staining. The tissues were fixed with ice-cold acetone for 20 min at -30 °C and dried at room temperature for 5 minutes followed with two times washing with Tris-buffered saline (TBS) containing 0.025% Triton X-100 (TBST). The sections were then incubated with background suppressor (23012A, Biotium) for 10 minutes and 10% normal goat serum (ab7481, Abcam) for two hours to block the non-specific staining. The primary anti-AR antibody was diluted in 1% bovine serum albumin (BSA) in TBST and incubated with the tissues at 4 °C for overnight, and the secondary antibody was incubated at 37 °C for one hour in dark. The tissues were counterstained with DAPI (5 μg/mL) in dark for 10 minutes, washed with TBST for three times and then mounted with ProLong Glass Antifade Mountant (P36982, Invitrogen).
